# *In silico* approach on drug repurposing - Antimalarial drugs against HIV-1 protease

**DOI:** 10.1101/2021.01.12.426148

**Authors:** Charli Deepak Arulanandam

## Abstract

Acquired Immunodeficiency Syndrome (AIDS), belonging to the retrovirus family is one of the most devastating contagious diseases of this century. Most of the available approved drugs are small molecules which are used in antiretroviral therapy (ART) that trigger the therapeutic response through binding to a targeted protein, HIV-1 protease (PR). This protein represents the most important antiretroviral drug target due to its key role in viral development inhibition. Computational tools using computer-aided technologies have proven useful in accelerating the drug discovery. In this study we evaluated selected FDA (USA) approved antimalarial drugs against HIV-1 protease to find a potential inhibitor candidate for HIV-1 PR (PDB 6DJ1). Binding affinities and Ki inhibition constant of an AutoDock 4.2 study suggest that of all assessed antimalarial agents, Lumefantrine (LUM) would be a most promising HIV-1 PR inhibitor.

## Introduction

Human Immunodeficiency Virus (HIV) produces dimeric aspartyl protease, which specifically cleaves the polyprotein precursors that encode structural proteins and enzymes of viruses (Debouck, 1992). HIV Protease (PR) has significant role in the viral life cycle (Chatterjee, Mridula, Mishra, Mittal and Hosur, 2005). Protease inhibitors (PIs) are vital drugs in active antiretroviral therapy for HIV-1 infections (Ellis *et al*., 2008). HIV-1 PR is an imperative drug target due to its key role in viral growth (Zondagh, Balakrishnan, Achilonu, Dirr and Sayed, 2018). Such proteolytic activity is vital for the production of matured virions and is, therefore, a striking target for therapeutic intervention (Stebbins and Debouck, 1997). Since both PR inhibitors (PIs) with reverse transcriptase inhibitors are included in the therapy, reductions in AIDS-associated deaths were observed in the mid-90s (Pau and George, 2014). HIV PIs, hustling drug resistance when mutations arise to the active and nonactive site of the PR (Nayak, Chandra, and Singh, 2019). However, long-term usage of these drugs induce mutations that cause drug resistance that challenge the long-term effectiveness in the medication of HIV-1-infected individuals (Havlir and Gandhi, 2015). A newer approach to anti-retroviral therapy is to design allosteric inhibitors against protease (Desai, Dikshit, and Iyer, 2012). Allosteric inhibitors are meanwhile studied extensively for developing therapeutics that prevent the proliferation of the virus. Certain antiretroviral agents constrain malaria-parasite growth, also some antimalarial drugs were shown to have weak antiretroviral effects (Skinner-Adams, McCarthy, Gardiner and Andrews, 2008). Computational techniques for binding-affinity prediction and molecular docking were considered in terms of their utility for drug discovery (Meng, Zhang, Mezei, and Cui, 2011). This study screens for appropriate inhibitors of this life-threatening viral enzyme. Most FDA-approved drugs prompt a therapeutic retort by binding to a target biological macromolecule (Feinberg *et al*., 2018). Computational techniques for binding-affinity prediction and molecular docking have long been utilized for the drug discovery (Leelananda and Lindert, 2016). AutoDock (AD) is appropriate for the docking studies for drug-like molecules binding to the viral protein targets (Forli *et al*., 2016). To screen for suitable antimalarials to treat or prevent HIV, we used a docking approach, which predicted the ligand-binding score and inhibition constant (Ki). The human Acquired Immunodeficiency Syndrome (AIDS) is caused by a lentivirus group belonging to the group of retroviruses. First identified in 1981, HIV is the cause of one of mankind’s fatal and most persistent epidemics. HIV produces retroviral aspartyl protease to cleave the structural proteins of viruses (Stingone, Sarmati, and Andreoni, 2020) and this proteolytic activity is required for viral maturation (Stebbins and Debouck, 1997). HIV protease (PR) shows an imperative role in the viral life cycle (Chatterjee *et al*., 2005). From this scenario, we know that protease inhibitors (PIs) are vital drugs to treat HIV-1 viral infections (Ellis *et al*., 2008). HIV-1 PR is a significant drug target due to its key role in viral growth (Arts and Hazuda, 2012; Zondagh *et al*., 2018). Because of the inclusion of reverse transcriptase (RT) with protease inhibitors (PIs) to reduce the AIDS-associated deaths were observed in the mid-90s (Pau and George, 2014). Antiretroviral chemotherapy fails if there is drug resistance. Medicines that once controlled HIV, may not be effective anymore if drug resistance evolves and spread. Drug-resistant HIV patients commonly have drug resistance to HIV treating drugs (https://aidsinfo.nih.gov). Hence, there is a constant demand for ever more effective anti-HIV compounds (Qureshi, Rajput, Kaur, and Kumar, 2018). There is currently no vaccine to prevent HIV infection. Hence, researchers are trying different approaches to find alternative solutions (https://www.hiv.gov). The most effective medicines among HIV PIs are Lopinavir (LPV) and Darunavir (DRV). However, they are fostering drug resistance due to mutations occurring either at the active or non-active site of the protein (Nayak *et al*., 2019). Particularly the continuous use of these drugs induces mutations leading to drug resistance, which challenge long-lasting efficacy in the medication of HIV-1-infected individuals (Havlir and Gandhi, 2015). For low-income patients drug resistance provides particularly a serious health threat (Pennings, 2013). Research in HIV is a challenging and intricate task that is highly resource-demanding (time-, money-, and manpower-consuming (Amzel et al., 2013). Drug-repositioning studies resulted in the pioneering of computational methods for the identification of highly suitable drug candidates from already approved ones (Jin & Wong, 2014). Such repositioned drugs would lessen the time and other efforts to develop promising drug candidates. These could go directly to pre-clinical testing and clinical trials, further reducing developing costs (Ashburn and Thor, 2004). Certain antiretroviral agents can inhibit the growth of the malaria-causing protist *Plasmodium*. Besides, some antimalarial drugs have weak antiretroviral effects (Skinner-Adams *et al*., 2008). Cheminformatic tools were found to be extremely valuable in medicinal drug development (Poli, Galati, Martinelli, Supuran, and Tuccinardi, 2020). The present study focuses on identifying lead compounds for this life-threatening viral enzyme from FDA-approved antimalarial drugs by using AutoDock (AD) 4.2. A therapeutic response via binding to a targeted enzyme is mainly achieved by small organic molecules - which holds for the majority of drugs with FDA-approval (Feinberg *et al*., 2018). Molecular docking pose and the prediction of binding-affinity are the main determinants for drug discovery by *in silico* approaches (Leelananda and Lindert, 2016). AD is suitable for docking studies for compounds that bind to proteins such as to HIV-1 PR (Forli *et al*., 2016). To predict the most suitable FDA approved antimalarial drug against HIV-1, we obtained the receptor–ligand binding score and the inhibitor constant Ki.

## Materials and Methods

### Target Protein

Global commitment to control the HIV/AIDS pandemic has increased (Meier, Evans, and Phelan, 2020). In this scenario AIDS-associated deaths reduced in the mid-90s because of the inclusion of both PR inhibitors (PIs) with reverse transcriptase inhibitors in therapy (Wong-Sam *et al*., 2018). The target protein is a Wild-type HIV-1 protease (PDB 6DJ1), protein resolution: 1.26 Å.

### Drug structural data

Antimalarial and HIV Drug data were retrieved from the PubChem database for this study (see Table 1 and Table 2). Structural data of drugs are available in a supplementary section. PubChem is an open repository for small molecules and their experimental biological activity (Bolton, Wang, Thiessen, and Bryant, 2008). It integrates and provides search, retrieval, visualization, analysis, and programatic access tools in an effort to exploit the efficacy of contributing information (Bolton *et al*., 2011).

### Cygwin

Cygwin was used to perform AutoDock 4.2 molecular docking in the Windows operating system (OS) (Rizvi, Shakil, and Haneef, 2013). Cygwin Packages are publicly available for docking operations at https://www.cygwin.com.

### AutoDock 4.2

AutoDock is an open-source application for performing molecular docking to predict ligand-receptor interactions (Pagadala, Syed, and Tuszynski, 2017). AutoDock is a serial application but several previous efforts parallel various aspects of the program (Norgan *et al*., 2011, 2011). In this study AutoDock 4.2 was used to predict the binding-affinity and molecular docking score for Wild-type HIV-1 PR (PDB 6DJ1)(Norgan *et al*., 2011). AD 4.2 suite is an open-source software for virtual screening and computational docking (Cosconati *et al*., 2010). This suite includes several complementary tools (Forli et al., 2016).

## 5. Computational methods

### 5.1. Targeted enzyme stability

In this study the target protein is a Wild-type HIV-1 protease (PDB 6DJ1) available from the RCSB PDB as a complex with Lopinavir (Rose *et al*., 2011) with a resolution of 1.26 Å (Wong-Sam *et al*., 2018). The native ligand (Lopinavir) water molecules were removed from PDB 6DJ1 for the docking study. The enzyme stability was calculated by using GROMACS molecular dynamics simulation (York *et al*., 1993; Lemkul, 2018) prior to the molecular docking experiment.

### 5.2. Ligands for docking study

Antimalarial and HIV Drug data were collected from PubChem for this study (see Table 1 and Table 2). PubChem is a public data repository that provides experimentally obtained data for small molecules (Bolton *et al*., 2008). PubChem allows the search for and retrieval as well as visualization of chemical information (Bolton *et al*., 2011).

### 5.3. Cygwin

The cygwin software was used to perform AutoDock 4.2 molecular docking in the Windows Operating System (OS)(Sharma *et al*., 2019). Cygwin is publicly available and free to download from: https://www.cygwin.com.

### 5.4. Molecular docking

In this molecular docking approach AD 4.2 was used to predict the binding-affinity, molecular docking pose, and Ki for the viral enzyme (PDB 6DJ1). AD 4.2 predicts receptor-ligand interactions and is publicly available for molecular docking studies (Pagadala *et al*., 2017). AD provides alternative functions of the program (Norgan *et al*., 2011) and this docking program is an open-source platform for virtual screening and computational docking (Cosconati *et al*., 2010).

#### 5.4.1. Grid generation

In this experiment, the grid box dimension values were attuned up to Grid center X-dimension: 100, Y-dimension: 126, Z-dimension: 126; Grid box spacing (in Ångström) X-center: 15.381, Y-center: 17.175, Z-center: 65.532 with an exhaustiveness value=8 (Abbasi *et al*., 2018). All the test compounds were docked distinctly with HIV-1 protease structure (PDB ID: 6DJ1). Generated docked enzyme inhibitor complex structures from the tool were assessed on the basis of minimal binding energy (kcal/mol) values and the patterns of hydrogen interaction visualized in BIOVIA Discovery Studio (DS) (Dassault Systèmes BIOVIA, 2016).

## 6. Docking and data visualization

All test compounds were individually docked with the X-ray crystallography (XRC) structure of HIV-1 protease (PDB ID: 6DJ1). The docked protein-ligand complexes were compared for interaction patterns of hydrogen and best value fits of binding energy (kcal/mol).

## Results and Discussion

Targeted enzymes were simulated in a water model prior to the molecular docking study using GROMACS to know its enzyme stability. Gyration of the test enzyme is shown in Fig. 1. This study targeted the identify of specific inhibitors of this critical viral enzyme from approved antimalarial drugs. Based on AutoDock 4.2 we revealed suitable HIV protease inhibitors from this source. Discovery studio was used to visualize the protein and docking results. AutoDock 4.2 based docking was used to approach aid to identify novel HIV protease inhibitors. AutoDock was carried out, which uncovers two lead molecules from a list of selected antimalarial drugs (see Table 1). From the tested drugs, we obtained the docking score for FDA approved antimalarial and anti-HIV drugs shown in Table 1 and STable 2. From the comparison of docking score and Ki inhibition, constant values show that the antimalarial drug LUM binding affinity was −13.19 kcal/mol, with an inhibition constant of 213.55 pM against HIV-1 protease. This lowest binding affinity and Ki value of LUM provided the best value compared to other tested ligands. Antimalarial drugs were docked *in silico* and the free binding energy and Ki values were obtained by using open source cygwin and AD tools. We found that LUM showed the best binding affinity and Ki values in virtual docking platform. Hence, we concluded that LUM has potency against HIV-1 PR. Antimalarial drugs docked to calculate the binding free energy to screen a suitable drug candidate against HIV-1 PR. The findings reported here provide therapeutic potentials against HIV-1. LUM was originally recommended for the treatment of uncomplicated malaria (https://www.who.int).

**Figure 1.**
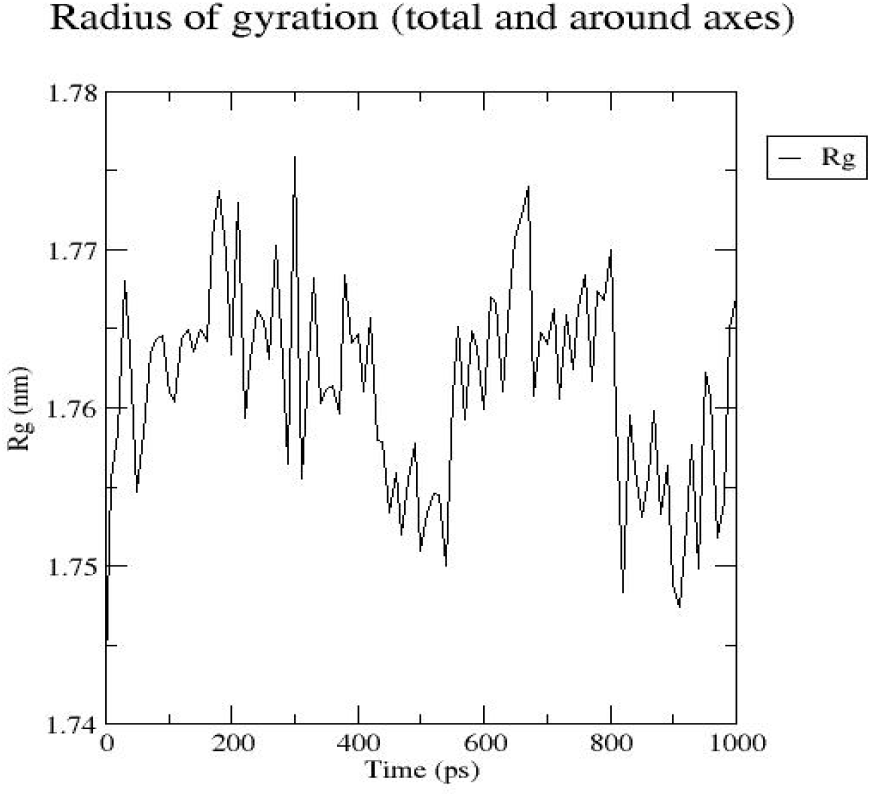
GROMACS: Gyration of targeted viral protein

## Conclusion

Molecular simulation and docking approaches were used to find a HIV protease inhibitor. We conclude from the obtained docking results using molecular docking tools that LUM might have potency to interrupt the HIV-1 maturation. To confirm LUM activity, we need to perform a Nucleic Acid Amplification Test (NAAT or NAT) in HIV-1 infected probands after treating them with LUM. Since this would be a clinical test later in drug approval for the repurposing of LUM as a HIV remedy.

## Supporting information

Supporting Inforation

## References

Abbasi MA, Hassan M, Aziz-ur-Rehman Siddiqui SZ, Shah SAA, Raza H, Seo S Y (2018). Synthesis, enzyme inhibitory kinetics mechanism and computational study of N -(4-methoxyphenethyl)-N -(substituted)-4-methylbenzenesulfonamides as novel therapeutic agents for Alzheimer’s disease. Journal of Life and Environmental Sciences, 6: e4962. DOI 10.7717/peerj.4962.

Amzel A, Toska E, Lovich R, Widyono M, Patel T, Foti C…, Altschuler J (2013). Promoting a Combination Approach to Paediatric HIV Psychosocial Support. AIDS, 27: S147–S157. DOI 10.1097/QAD.0000000000000098.

Arts EJ, Hazuda DJ (2012). HIV-1 Antiretroviral Drug Therapy. Cold Spring Harbor Perspectives in Medicine, 2: a007161– a007161. DOI 10.1101/cshperspect.a007161.

Ashburn T T, Thor KB (2004). Drug repositioning: identifying and developing new uses for existing drugs. Nature Reviews Drug Discovery, 3: 673–683. DOI 10.1038/nrd1468.

Bolton E E, Chen J, Kim, S, Han L, He S, Shi W, Bryant S H (2011). PubChem3D: A new resource for scientists. Journal of Cheminformatics, 3: 32. DOI 10.1186/1758-2946-3-32.

Bolton E E, Wang Y, Thiessen P A, Bryant S H (2008). Chapter 12 PubChem: Integrated Platform of Small Molecules and Biological Activities. In Annual Reports in Computational Chemistry 4: 217–241. DOI 10.1016/S1574-1400(08)00012-1.

Chatterjee A, Mridula P, Mishra RK, Mittal R, Hosur RV (2005). Folding regulates autoprocessing of HIV-1 protease precursor. Journal of Biological Chemistry, 280: 11369– 11378. DOI 10.1074/jbc.M412603200.

Cosconati S, Forli S, Perryman AL, Harris R, Goodsell DS, Olson AJ (2010). Virtual screening with AutoDock: theory and practice. Expert Opinion on Drug Discovery, 5: 597–607. DOI 10.1517/17460441.2010.484460.

Debouck C (1992). The HIV-1 Protease as a Therapeutic Target for AIDS. AIDS Research and Human Retroviruses, 8: 153–164. DOI 10.1089/aid.1992.8.153.

Desai M, Dikshit R, Iyer G (2012). Antiretroviral drugs: Critical issues and recent advances. Indian Journal of Pharmacology, 44: 288. DOI 10.4103/0253-7613.96296.

Ellis RJ, Marquie-Beck J, Delaney P, Alexander T, Clifford DB, McArthur JC, Grant I (2008). Human immunodeficiency virus protease inhibitors and risk for peripheral neuropathy. Annals of Neurology, 64: 566–572. DOI 10.1002/ana.21484.

Feinberg EN, Sur D, Wu Z, Husic BE, Mai H, Li Y, Pande VS (2018). PotentialNet for Molecular Property Prediction. ACS Central Science, 4: 1520–1530. DOI 10.1021/acscentsci.8b00507.

Forli S, Huey R, Pique M, Sanner MF, Goodsell DS, Olson AJ (2016). Computational protein–ligand docking and virtual drug screening with the AutoDock suite. Nature Protocols, 11: 905–919. DOI 10.1038/nprot.2016.051.

Havlir D, Gandhi M (2015). Implementation challenges for longacting antivirals as treatment. Current Opinion in HIV and AIDS 10: 282–289. DOI 10.1097/COH.0000000000000158.

Jin G, Wong STC (2014). Toward better drug repositioning: prioritizing and integrating existing methods into efficient pipelines. Drug Discovery Today 19: 637–644. DOI 10.1016/j.drudis.2013.11.005.

Leelananda SP, Lindert S (2016). Computational methods in drug discovery. Beilstein Journal of Organic Chemistry 12: 2694– 2718. DOI 10.3762/bjoc.12.267.

Meier BM, Evans DP, Phelan A (2020). Rights-Based Approaches to Preventing, Detecting, and Responding to Infectious Disease Outbreaks. SSRN Electronic Journal 217–253. DOI 10.2139/ssrn.3560669.

Meng XY, Zhang HX, Mezei M, Cui M (2011). Molecular Docking: A Powerful Approach for Structure-Based Drug Discovery. Current Computer Aided-Drug Design 7: 146–157. DOI 10.2174/157340911795677602.

Nayak C, Chandra I, Singh SK (2019). An in silico pharmacological approach toward the discovery of potent inhibitors to combat drug resistance HIV-1 protease variants. Journal of Cellular Biochemistry 120: 9063–9081. DOI 10.1002/jcb.28181.

Norgan AP, Coffman PK, Kocher JPA, Katzmann DJ, Sosa CP (2011). Multilevel Parallelization of AutoDock 4.2. Journal of Cheminformatics 3: 12. DOI 10.1186/1758-2946-3-12.

Pagadala NS, Syed K, Tuszynski J (2017). Software for molecular docking: a review. Biophysical Reviews 9: 91–102. DOI 10.1007/s12551-016-0247-1.

Pau AK, George JM (2014). Antiretroviral Therapy. Infectious Disease Clinics of North America 28: 371–402. DOI 10.1016/j.idc.2014.06.001

Pennings PS (2013). HIV drug resistance: problems and perspectives. Infectious Disease Reports 5: 5. DOI 10.4081/idr.2013.s1.e5.

Poli G, Galati S, Martinelli A, Supuran CT, Tuccinardi T (2020). Development of a cheminformatics platform for selectivity analyses of carbonic anhydrase inhibitors. Journal of Enzyme Inhibition and Medicinal Chemistry 35: 365–371. DOI 10.1080/14756366.2019.1705291.

Qureshi A, Rajput A, Kaur G, Kumar M (2018). HIVprotI: an integrated web based platform for prediction and design of HIV proteins inhibitors. Journal of Cheminformatics 10: 12. DOI 10.1186/s13321-018-0266-y.

Rizvi SMD, Shakil S, Haneef M (2013). A simple click by click protocol to perform docking: AutoDock 4.2 made easy for non-bioinformaticians. Excli Journal 12: 831.

Rose PW, Beran B, Bi C, Bluhm WF, Dimitropoulos D, Goodsell DS, Bourne PE (2011). The RCSB Protein Data Bank: redesigned web site and web services. Nucleic Acids Research 39: D392– D401. DOI 10.1093/nar/gkq1021.

Sharma A, Islam MH, Fatima N, Upadhyay TK, Khan MKA, Dwivedi UN, Sharma R (2019). Elucidation of marine fungi derived anthraquinones as mycobacterial mycolic acid synthesis inhibitors: an in silico approach. Molecular Biology Reports 46: 1715–1725. DOI 10.1007/s11033-019-04621-0.

Skinner-Adams TS, McCarthy JS, Gardiner DL, Andrews KT (2008). HIV and malaria co-infection: interactions and consequences of chemotherapy. Trends in Parasitology 24: 264–271. DOI 10.1016/j.pt.2008.03.008.

Stebbins J, Debouck C (1997). A Microtiter Colorimetric Assay for the HIV-1 Protease. Analytical Biochemistry 248: 246–250. DOI 10.1006/abio.1997.2111.

Stingone C, Sarmati L, Andreoni M (2020). The Clinical Spectrum of Human Immunodeficiency Virus Infection. In: Sexually Transmitted Infections 295–317. DOI 10.1007/978-3-030-02200-6_15.

Wong-Sam A, Wang YF, Zhang Y, Ghosh AK, Harrison RW, Weber IT (2018). Drug Resistance Mutation L76V Alters Nonpolar Interactions at the Flap–Core Interface of HIV-1 Protease. ACS Omega 3: 12132–12140. DOI 10.1021/acsomega.8b01683.

Zondagh J, Balakrishnan V, Achilonu I, Dirr HW S ayed Y (2018). Molecular dynamics and ligand docking of a hinge region variant of South African HIV-1 subtype C protease. Journal of Molecular Graphics and Modelling, 82: 1–11. DOI 10.1016/j.jmgm.2018.03.006.

York DM, Darden TA, Pedersen LG, Anderson MW. Molecular dynamics simulation of HIV-1 protease in a crystalline environment and in solution. Biochemistry. 1993 Feb 1;32(6):1443–53.

Lemkul J. From proteins to perturbed Hamiltonians: A suite of tutorials for the GROMACS-2018 molecular simulation package [article v1. 0]. Living Journal of Computational Molecular Science. 2018 Oct 27;1(1):5068.

