## Supporting Inforation for "*In silico* approach on drug repurposing - Antimalarial drugs against HIV-1 protease"

Charli Deepak Arulanandam*

Department of Medicinal and Applied Chemistry, Kaohsiung Medical University, Kaohsiung, Taiwan

**Contents:** Figures and Tables regarding information of utilized molecules

**Figure section**

**Figure.S.1. Wild-type HIV-1 protease as a drug target**

**Figure.S.2. Cavity of active site of wild-type HIV-1 protease.**

**Figure.S.3. Interaction of Native ligand-Lopinavir with HIV-1 protease.**

**Figure S.4. Antimalarials as test compounds.**

**Figure S.5. Anti-HIV drugs as test compounds**

**Table section**

**Table. S.1. Available HIV medicines and drug targets**

### Background study for the test compounds and HIV viral protein selection

Treatment with HIV medicines is known by the name of antiretroviral therapy (ART). A person's initial HIV regimen generally includes three HIV medicines from at least two different drug classes. ART can't cure HIV, but HIV medicines help people with HIV live longer, healthier lives. HIV medicines also reduce the risk of HIV transmission. HIV patients prone to other infectious disease. In Africa and other developing countries HIV + malaria increases the death rate of the HIV people. Table.S1. lists the HIV medicines recommended for the treatment of HIV infection. Here the medicines are listed according to drug class and identified by the generic and brand names. Available approved drug data information collected from the AIDSinfo. It is accessible from the website www.aidsinfo.nih.gov. HIV PR achieves a crucial role in viral replication by processing the viral precursor polyproteins into mature viral proteins. Inhibitors bind in the active-site cavity of dimeric HIV PR and block its catalytic activity. More than 100 mutations in the PR gene have been associated with drug resistance (Rhee et al., 2016). As per literature search a decrease in AIDS-associated deaths was observed in the mid-90s because of the inclusion of both PR inhibitors (PIs) with reverse transcriptase inhibitors in therapy (Konvalinka et al., 2015).

### Selection of test ligand for drug repurposing

HIV and malaria are the two most prevalent and deadly diseases in the world. This co-infection accounted for about 255 million cases in 2017 (Del-Tejo et al., 2020). These common infections in Africa causes the substantial morbidity and mortality in prenatal women. There was high prevalence of malaria and HIV mortality among severely undernourished children (Jacques et al., 2020). "Co-infection" differs from single infection mainly in the effect of co-infection on the host health which may be harmful, beneficial or absent. This effect occurs mainly due to changes in host immune response (Abbas e al., 2020). A robust and effective malaria and HIV control management program should be strongly underpinned (Obioma et al., 2020). To resolve co-infection in this study we choose antimalarial drug candidate to evaluate against the HIV-1 protease.

### Target Protein Preparation

X-ray crystallographic structure accessed from the RCSB protein data bank and the structural data retrieved as a PDB file. This protein 3D structure comprises native ligand and water molecules. So, first we removed the water and native ligand from the PDB data file. To do this GROMACS used by using below commands.

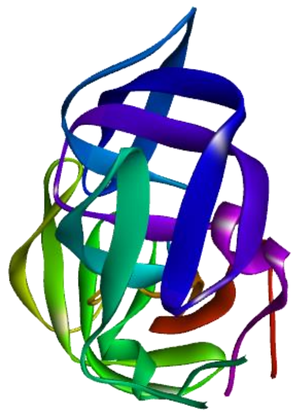

**Figure.S.1.** Wild-type HIV-1 protease as a drug target [PDB ID: 6DJ1- Native ligand and water molecules removed from the target protein].

### Automated Version of Active Site Prediction (AADS)

Active site of HIV-1 enzyme predicted by using AADS web server. This server available from the web link http://www.scfbio-iitd.res.in/dock/ActiveSite_new.jsp

Predicted active site has 195 amino acids.

MKFKMFKIGTRIGPGLDLDREIRIPGPTDDRLREIVGVPEPGTGGKRRDLDLIVIGPTIGPWNRRRDRRDLLEGNGIIINKTDRDDDLD
DVVEIIWNKRRRLLVLVAINIINRADAVGLINITIIGITRPAGLALIPIIGIGLAIAGIIPGLIPGGLDIDIGIIAVPDGVATLGAVGVATGLVD
DTVRTLRDRDR

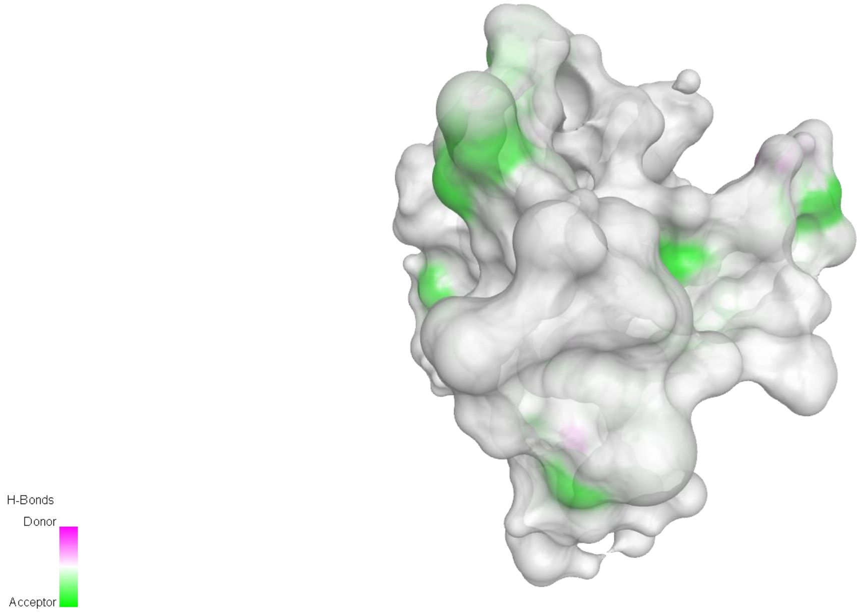


**Figure.S.2. Cavity of active site of wild-type HIV-1 protease.**

**Molecular Docking study:**

AD 4.2 based docking was used in this work to identify the novel HIV-1 protease inhibitor. Discovery studio was used to visualize the protein and docking results. Docking study to uncovers the lead molecules from a list of selected antimalarial drugs. We performed this molecular docking study based on a simple protocol from the Rizvi et al., 2013.

### 2D diagram of native ligand interaction

Unfavorable bump noted from the analysis of 2D diagram of native ligand (LPV) and interface of the viral enzyme. Unfavorable bump in molecular docking does not necessarily mean that compound is not a good inhibitor for many reasons. For example, LPV drug might have an unexpected mode of action or a different binding site.

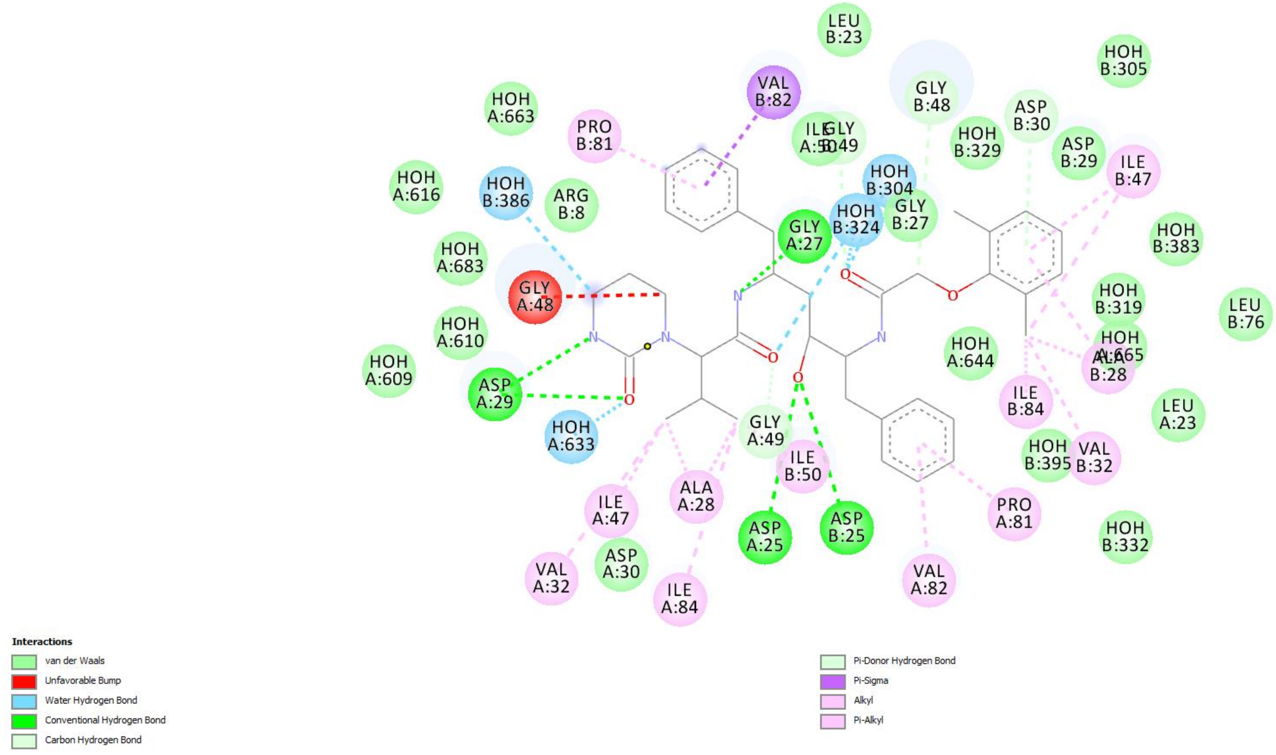


**Figure.S.3. Interaction of Native ligand-Lopinavir with HIV-1 protease.**

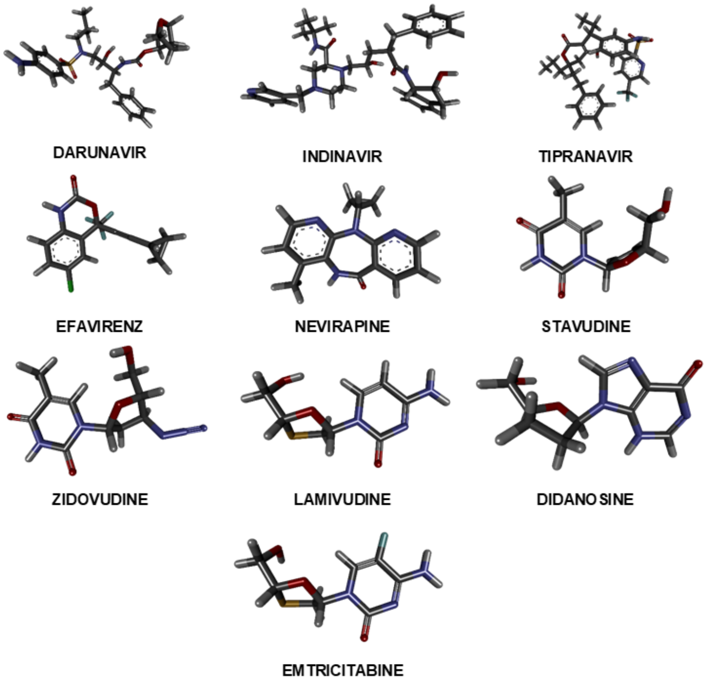


**Figure S.4. Antimalarials as test compunds** [3D- Ball and Stick models].

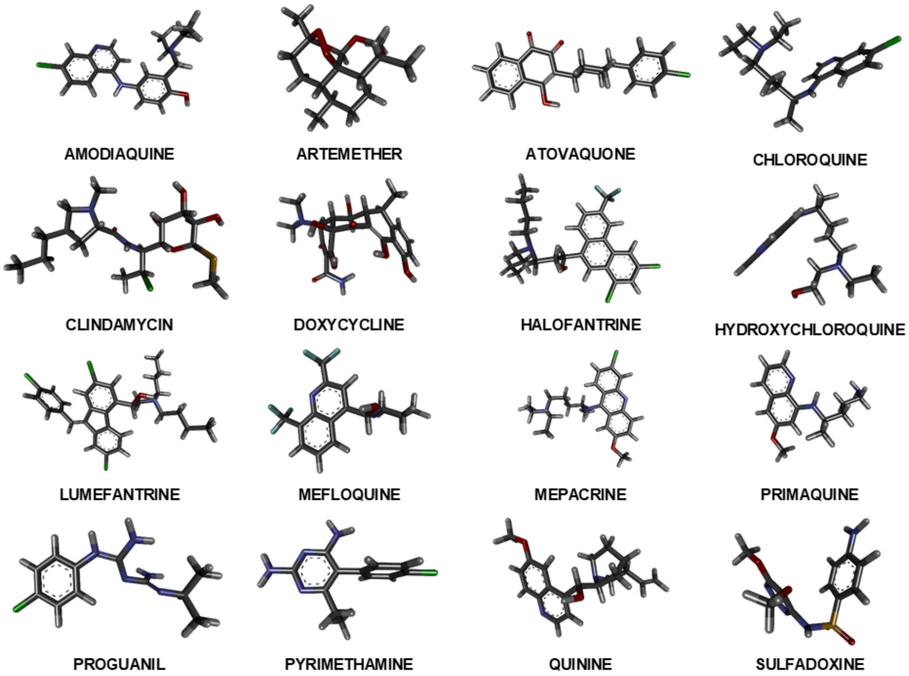


**Figure S.5. Anti-HIV drugs as test compounds** [3D - Ball and Stick models].

### GROMACS

GROningen MAchine for Chemical Simulations (GROMACS) is a molecular dynamics simulation (MDS) package intended for the simulations of biological macromolecules. It was initially developed in the Biophysical Chemistry department of University of Groningen. It is one of the fastest and most popular software packages for the MDS. This open-source software can run on graphics processing units (GPUs) and central processing units (CPUs).

### Molecular dynamics simulation

Following commends used for the process of setting up a simulation system containing a viral protein (6dj1) in a virtual box of water, with ions. This methodology established based on the information available from http://www.mdtutorials.com/gmx/lysozyme/index.html

To open the working Directory:

cd /

cd gromacs-2020.3

cd build

*Step 1: Prepare the Topology*

gmx pdb2gmx -f 6dj1_clean.pdb -o 6dj1_processed.gro -water spce

choose the force filed, we choose for this study OPLS-AA/L all-atom force field

*Step 2: Defining the Unit Cell & Adding Solvent*

gmx editconf -f 6dj1_processed.gro -o 6dj1_newbox.gro -c -d 1.0 -bt cubic

gmx solvate -cp 6dj1_newbox.gro -cs spc216.gro -o 6dj1_solv.gro -p topol.top

*Step 3: Addding Ions*

gmx grompp -f ions.mdp -c 6dj1_solv.gro -p topol.top -o ions.tpr

gmx genion -s ions.tpr -o 6dj1_solv_ions.gro -p topol.top -pname NA -nname CL –neutral

Type Nano ions.mdp to save the parameters in LINUX terminal and paste the ions.mdp parameters to create ions.mdp file. We used ions.mdb parameters according to the GROMACS tutorial which is given at the end of this supporting file.

*Step 4: Energy Minimization*

gmx grompp -f minim.mdp -c 6dj1_solv_ions.gro -p topol.top -o em.tpr

gmx mdrun -v -deffnm em

*Step 5: NVT Equilibration*

gmx grompp -f nvt.mdp -c em.gro -r em.gro -p topol.top -o nvt.tpr

gmx mdrun -deffnm nvt

*Step 6: NPT Equilibration*

gmx grompp -f npt.mdp -c nvt.gro -r nvt.gro -t nvt.cpt -p topol.top -o npt.tpr

gmx mdrun -deffnm npt

*Step 7: Production MD*

gmx grompp -f md.mdp -c npt.gro -t npt.cpt -p topol.top -o md_0_1.tpr

gmx mdrun -deffnm md_0_1

*Step 7: Analysis*

gmx trjconv -s md_0_1.tpr -f md_0_1.xtc -o md_0_1_noPBC.xtc -pbc mol –center

gmx gyrate -s md_0_1.tpr -f md_0_1_noPBC.xtc -o gyrate.xvg

### Parameters for the GROMACS Molecular dynamics simulation

### ions.mdp

```
; ions.mdp - used as input into grompp to generate ions.tpr
; Parameters describing what to do, when to stop and what to save
integrator  = steep         ; Algorithm (steep = steepest descent minimization)
emtol       = 1000.0        ; Stop minimization when the maximum force < 1000.0 kJ/mol/nm
emstep      = 0.01          ; Minimization step size
nsteps      = 50000         ; Maximum number of (minimization) steps to perform

; Parameters describing how to find the neighbors of each atom and how to calculate the interactions
nstlist         = 1         ; Frequency to update the neighbor list and long range forces
cutoff-scheme             = Verlet    ; Buffered neighbor searching
ns_type         = grid      ; Method to determine neighbor list (simple, grid)
coulombtype     = cutoff    ; Treatment of long range electrostatic interactions
rcoulomb        = 1.0       ; Short-range electrostatic cut-off
rvdw            = 1.0       ; Short-range Van der Waals cut-off
pbc             = xyz       ; Periodic Boundary Conditions in all 3 dimensions
```

### minim.mdp

```
; minim.mdp - used as input into grompp to generate em.tpr
; Parameters describing what to do, when to stop and what to save
integrator  = steep         ; Algorithm (steep = steepest descent minimization)
emtol       = 1000.0        ; Stop minimization when the maximum force < 1000.0 kJ/mol/nm
emstep      = 0.01          ; Minimization step size
nsteps      = 50000         ; Maximum number of (minimization) steps to perform

; Parameters describing how to find the neighbors of each atom and how to calculate the interactions
nstlist         = 1         ; Frequency to update the neighbor list and long range forces
cutoff-scheme   = Verlet    ; Buffered neighbor searching
ns_type         = grid      ; Method to determine neighbor list (simple, grid)
coulombtype     = PME       ; Treatment of long range electrostatic interactions
rcoulomb        = 1.0       ; Short-range electrostatic cut-off
rvdw            = 1.0       ; Short-range Van der Waals cut-off
pbc             = xyz       ; Periodic Boundary Conditions in all 3 dimensions
```

### nvt.mdp

```
title                = OPLS Lysozyme NVT equilibration
define               = -DPOSRES  ; position restrain the protein
; Run parameters
integrator           = md        ; leap-frog integrator
nsteps               = 50000     ; 2 * 50000 = 100 ps
dt                   = 0.002     ; 2 fs
; Output control
nstxout              = 500       ; save coordinates every 1.0 ps
nstvout              = 500       ; save velocities every 1.0 ps
nstenergy            = 500       ; save energies every 1.0 ps
nstlog               = 500       ; update log file every 1.0 ps
; Bond parameters
continuation         = no        ; first dynamics run
constraint_algorithm    = lincs     ; holonomic constraints
constraints          = h-bonds   ; bonds involving H are constrained
lincs_iter           = 1         ; accuracy of LINCS
lincs_order          = 4         ; also related to accuracy
; Nonbonded settings
cutoff-scheme        = Verlet    ; Buffered neighbor searching
ns_type              = grid      ; search neighboring grid cells
nstlist              = 10        ; 20 fs, largely irrelevant with Verlet
rcoulomb             = 1.0       ; short-range electrostatic cutoff (in nm)
rvdw                 = 1.0       ; short-range van der Waals cutoff (in nm)
DispCorr             = EnerPres  ; account for cut-off vdW scheme
; Electrostatics
coulombtype          = PME       ; Particle Mesh Ewald for long-range electrostatics
pme_order            = 4         ; cubic interpolation
fourierspacing       = 0.16      ; grid spacing for FFT
; Temperature coupling is on
tcoupl               = V-rescale             ; modified Berendsen thermostat
tc-grps              = Protein Non-Protein   ; two coupling groups - more accurate
tau_t                = 0.1     0.1           ; time constant, in ps
ref_t                = 300     300           ; reference temperature, one for each group, in K
; Pressure coupling is off
pcoupl               = no        ; no pressure coupling in NVT
; Periodic boundary conditions
pbc                  = xyz       ; 3-D PBC
; Velocity generation
gen_vel              = yes       ; assign velocities from Maxwell distribution
gen_temp             = 300       ; temperature for Maxwell distribution
gen_seed             = -1        ; generate a random seed
```

### npt.mdp

```
title                = OPLS Lysozyme NPT equilibration
define               = -DPOSRES  ; position restrain the protein
; Run parameters
integrator           = md        ; leap-frog integrator
nsteps               = 50000     ; 2 * 50000 = 100 ps
dt                   = 0.002     ; 2 fs
```

```
; Output control
nstxout          = 500    ; save coordinates every 1.0 ps
nstvout          = 500    ; save velocities every 1.0 ps
nstenergy        = 500    ; save energies every 1.0 ps
nstlog           = 500    ; update log file every 1.0 ps
; Bond parameters
continuation         = yes     ; Restarting after NVT
constraint_algorithm = lincs   ; holonomic constraints
constraints          = h-bonds ; bonds involving H are constrained
lincs_iter           = 1       ; accuracy of LINCS
lincs_order          = 4       ; also related to accuracy
; Nonbonded settings
cutoff-scheme    = Verlet    ; Buffered neighbor searching
ns_type          = grid      ; search neighboring grid cells
nstlist          = 10        ; 20 fs, largely irrelevant with Verlet scheme
rcoulomb         = 1.0       ; short-range electrostatic cutoff (in nm)
rvdw             = 1.0       ; short-range van der Waals cutoff (in nm)
DispCorr         = EnerPres  ; account for cut-off vdW scheme
; Electrostatics
coulombtype      = PME       ; Particle Mesh Ewald for long-range electrostatics
pme_order        = 4         ; cubic interpolation
fourierspacing   = 0.16      ; grid spacing for FFT
; Temperature coupling is on
tcoupl           = V-rescale            ; modified Berendsen thermostat
tc-grps          = Protein Non-Protein  ; two coupling groups - more accurate
tau_t            = 0.1   0.1            ; time constant, in ps
ref_t            = 300   300            ; reference temperature, one for each group, in K
; Pressure coupling is on
pcoupl           = Parrinello-Rahman    ; Pressure coupling on in NPT
pcoupltype       = isotropic            ; uniform scaling of box vectors
tau_p            = 2.0                  ; time constant, in ps
ref_p            = 1.0                  ; reference pressure, in bar
compressibility  = 4.5e-5               ; isothermal compressibility of water, bar^-1
refcoord_scaling = com
; Periodic boundary conditions
pbc              = xyz       ; 3-D PBC
; Velocity generation
gen_vel          = no        ; Velocity generation is off
```

### md.mdp

```
title            = OPLS Lysozyme NPT equilibration
; Run parameters
integrator       = md        ; leap-frog integrator
nsteps           = 500000    ; 2 * 500000 = 1000 ps (1 ns)
dt               = 0.002     ; 2 fs
; Output control
nstxout          = 0         ; suppress bulky .trr file by specifying
nstvout          = 0         ; 0 for output frequency of nstxout,
nstfout          = 0         ; nstvout, and nstfout
nstenergy        = 5000      ; save energies every 10.0 ps
nstlog           = 5000      ; update log file every 10.0 ps
nstxout-compressed  = 5000   ; save compressed coordinates every 10.0 ps
compressed-x-grps   = System ; save the whole system
; Bond parameters
continuation     = yes       ; Restarting after NPT
```

```
constraint_algorithm    = lincs     ; holonomic constraints
constraints             = h-bonds   ; bonds involving H are constrained
lincs_iter              = 1         ; accuracy of LINCS
lincs_order             = 4         ; also related to accuracy
; Neighborsearching
cutoff-scheme           = Verlet    ; Buffered neighbor searching
ns_type                 = grid      ; search neighboring grid cells
nstlist                 = 10        ; 20 fs, largely irrelevant with Verlet scheme
rcoulomb                = 1.0       ; short-range electrostatic cutoff (in nm)
rvdw                    = 1.0       ; short-range van der Waals cutoff (in nm)
; Electrostatics
coulombtype             = PME       ; Particle Mesh Ewald for long-range electrostatics
pme_order               = 4         ; cubic interpolation
fourierspacing          = 0.16      ; grid spacing for FFT
; Temperature coupling is on
tcoupl                  = V-rescale             ; modified Berendsen thermostat
tc-grps                 = Protein Non-Protein   ; two coupling groups - more accurate
tau_t                   = 0.1     0.1           ; time constant, in ps
ref_t                   = 300     300           ; reference temperature, one for each group, in K
; Pressure coupling is on
pcoupl                  = Parrinello-Rahman     ; Pressure coupling on in NPT
pcoupltype              = isotropic             ; uniform scaling of box vectors
tau_p                   = 2.0                   ; time constant, in ps
ref_p                   = 1.0                   ; reference pressure, in bar
compressibility         = 4.5e-5                ; isothermal compressibility of water, bar^-1
; Periodic boundary conditions
pbc                     = xyz       ; 3-D PBC
; Dispersion correction
DispCorr                = EnerPres  ; account for cut-off vdW scheme
; Velocity generation
gen_vel                 = no        ; Velocity generation is off
```

**Table. S.1.** Available HIV medicines and drug targets.

| Chemical name | PubChem CID/SID | Commercial name |
|---|---|---|
| **Nucleoside Reverse Transcriptase Inhibitors (NRTIs):**<br>NRTIs block reverse transcriptase, an enzyme for the HIV replication. | | |
| abacavir | CID 441300 | Ziagen |
| emtricitabine | CID 60877 | Emtriva |
| lamivudine | CID 60825 | Epivir |
| tenofovir disoproxil fumarate | CID 6398764 | Viread |
| zidovudine | CID 35370 | Retrovir |
| **Non-Nucleoside Reverse Transcriptase Inhibitors (NNRTIs):** | | |
| doravirine | CID 58460047 | Pifeltro |
| efavirenz | CID 64139 | Sustiva |
| etravirine | CID 193962 | Intelence |
| nevirapine | CID 4463 | Viramune |
| rilpivirine | CID 6451164 | Edurant |

| | | |
|---|---|---|
| **Protease Inhibitors (PIs):**<br>PIs block HIV protease, an enzyme involving in HIV replication. | | |
| atazanavir | CID 148192 | Reyataz |
| darunavir | CID 213039 | Prezista |
| fosamprenavir | CID 131536 | Lexiva |
| ritonavir | CID 392622 | Norvir |
| saquinavir | CID 441243 | Invirase |
| tipranavir | CID 54682461 | Aptivus |
| **Fusion Inhibitors:** | | |
| enfuvirtide | CID 24847866 | Fuzeon |
| **CCR5 Antagonists:**<br>CCR5 antagonists block CCR5 co-receptors on the surface of certain immune cells that HIV needs to enter the cells. | | |
| maraviroc | CID 3002977 | Selzentry |
| **Integrase Inhibitors:**<br>Integrase inhibitors block HIV integrase, an enzyme which is involving in the replication of HIV. | | |
| dolutegravir | CID 54726191 | Tivicay |
| raltegravir | CID 54671008 | Isentress |
| **Post-Attachment Inhibitors:**<br>Post-attachment inhibitors block CD4 receptors on the surface of certain immune cells that HIV needs to enter the cells. | | |
| ibalizumab-uiyk | CID 160671956 | Trogarzo |
| **Pharmacokinetic Enhancers:**<br>Pharmacokinetic enhancers are used in HIV treatment to increase the effectiveness of an HIV medicine included in an HIV regimen. | | |
| cobicistat | CID 25151504 | Tybost |
| **Combination HIV Medicines:**<br>Combination HIV medicines contain two or more HIV medicines from one or more drug classes. | | |
| abacavir and lamivudine | CID 469584 | Epzicom |
| abacavir, dolutegravir, and lamivudine | CID 54736666 | Triumeq |
| abacavir, lamivudine, and zidovudine | CID 5726 | Trizivir |
| atazanavir and cobicistat | CID 86583336 | Evotaz |
| bictegravir, emtricitabine, and tenofovir alafenamide | SID 354339134 | Biktarvy |
| darunavir and cobicistat | CID 57327017 | Prezcobix |
| darunavir, cobicistat, emtricitabine, and tenofovir alafenamide | SID 384585360 | Symtuza |
| dolutegravir and lamivudine | SID 384585498 | Dovato |
| dolutegravir and rilpivirine | CID 131801472 | Juluca |
| doravirine, lamivudine, and tenofovir disoproxil fumarate | SID 384585374 | Delstrigo |

| | | |
|---|---|---|
| efavirenz, emtricitabine, and tenofovir disoproxil fumarate | CID 9833525 | Atripla |
| efavirenz, lamivudine, and tenofovir disoproxil fumarate | SID 384585370 | Symfi |
| efavirenz, lamivudine, and tenofovir disoproxil fumarate | | Symfi Lo |
| elvitegravir, cobicistat, emtricitabine, and tenofovir alafenamide | CID 66545969 | Genvoya |
| elvitegravir, cobicistat, emtricitabine, and tenofovir disoproxil fumarate | CID 44232548 | Stribild |
| emtricitabine, rilpivirine, and tenofovir alafenamide | CID 91810705 | Odefsey |
| emtricitabine, rilpivirine, and tenofovir disoproxil fumarate | CID 44128157 | Complera |
| emtricitabine and tenofovir alafenamide | CID 90469070 | Descovy |
| emtricitabine and tenofovir disoproxil fumarate | CID 464205 | Truvada |
| lamivudine and tenofovir disoproxil fumarate | SID 384585373 | Cimduo |
| lamivudine and zidovudine | CID 72187 | Combivir |
| lopinavir and ritonavir | CID 11979606 | Kaletra |

### References


Abbas AM, Salah S, Fathy SK, Rashad A, AboBakr A, Yousof EA, Saeed A, Youssef EN, Shaltout AS, Ahmed OA. Role of Co-Infection in the Immunopathology of COVID-19 in pregnancy.

Del-Tejo PL, Cubas-Vega N, Caraballo-Guerra C, da Silva BM, da Silva Valente J, Sampaio VS, Baia DC, Castro DB, Martinez-Espinosa FE, Siqueira AM, Lacerda MV. Should We Care About Plasmodium Vivax and HIV Coinfection? A Systematic Review and a Cases Series From the Brazilian Amazon.

Jacques M, Salissou MM, Kaswiyi L, Guan F, Lei J. Co-infection of malaria and HIV infection in severely undernourished children in the Democratic Republic of the Congo: a cross-sectional study. Parasitology. 2020 Feb;147(2):248-53.

Konvalinka J, Kräusslich HG, Müller B. Retroviral proteases and their roles in virion maturation. Virology. 2015 May 1;479:403-17.

Obioma A, Nnnena I, Judith N. Invivo Impact of Malaria and HIV Co-infection On CD4 Cell Count of Infected Patients of Niger Delta Extraction. Journal of Advanced Pharmaceutical Science and Technology. 2020 May 8;2(2):18.

Rhee SY, Sankaran K, Varghese V, Winters MA, Hurt CB, Eron JJ, Parkin N, Holmes SP, Holodniy M, Shafer RW. HIV-1 protease, reverse transcriptase, and integrase variation. Journal of virology. 2016 Jul 1;90(13):6058-70.

Rizvi SMD, Shakil S, Haneef M. A simple click by click protocol to perform docking: Autodock 4.2 made easy for non-bioinformaticians. EXCLI J 2013.
